# Biological consequences of asymmetric hop diffusion in the cell membrane

**DOI:** 10.1101/2025.06.04.657947

**Authors:** Damien Hall

## Abstract

Over the last 30 years, the hop diffusion model has been an important paradigm for interpreting both cell membrane structure and dynamics. The basic premise of the model is that the cell membrane is organized into small domain regions through a combination of multi-component phase-separation and interaction between membrane components and the proximal fibers of the intracellular cytoskeleton and extracellular matrix. The partitioned characteristics of this two-dimensional fluid are thought to impose both steric and hydrodynamic barriers that restrict the free motion of mobile membrane components, save for their occasional passage via ‘hopping’ from one domain to another. Previous investigations of hop diffusion within the cell membrane have identified the potential for diffusional anisotropy **[Jaqaman et al. 2011. Cell, 146(4), pp.593-606]**. This work utilizes numerical simulations and develops new analytical theory to provide an approximate quantitative description of such asymmetric compartmentalization. These methods are then used to examine the physical requirements for generation of asymmetric hop diffusion within the membrane before concluding with a discussion of the potential biological consequences of such behavior.

## Introduction

In most eukaryotic cells an external phospholipid membrane bilayer lies atop of the cortical cytoskeleton, which is typically composed of a dense meshwork of actin fibers and spectrin proteins **[Jacobson et al. 2019; Kalappurakkal et al. 2020]**. Connections between cell membrane and the cortical cytoskeleton are established by a range of linkers, of which the most prominent are the ERM (exrin, radaxin and moesin), catenin and ankyrin proteins **[Charras et al. 2006; Fehon et al. 2010]**. Depending on the type of cell and its position within a tissue, a second fibrous network called the extracellular matrix (ECM)^1^ can exist external to the membrane with ECM-membrane linkages carried out through laminin and integrin proteins **[Luna and Hitt, 1992; Theocharis et al. 2016; Changede et al. 2019]**. Linear persistence of the proximal fibers in the cortical cytoskeleton and ECM impart a similar linear ordering to the membrane proteins attached to them **[Jacobson et al. 2019; Kalappurakkal et al. 2019]**. These ordered arrays of stationary integral membrane proteins within the cell membrane are thought to create both short-range steric, and longer-range hydrodynamic, barriers to the movement of un-attached lipids and proteins existing in the phospholipid bilayer **(Fig. 1a)**. This conceptualization forms the basis of the hop diffusion model of the cell membrane **[Edelman, 1976; Sheetz et al. 1980; Koppel et al. 1981; Tsuji and Ohnishi, 1986; Saxton, 1990; Sako and Kusumi, 1994; Saxton and Jacobson, 1997; Kusumi et al. 2005; Kalappurakkal et al. 2020]** which provides a reasonable physical explanation for the observed motion of individual mobile membrane components as they undergo repeated frustrated diffusion within a bounded region with the occasional rare ‘hop’ (escape) across a barrier to enter a neighboring compartment **[Andrade et al. 2015;Goiko et al 2016; Sadegh et al. 2017; de Wit et al, 2018; Mashanov et al. 2021; van der Linden, 2022]**.

**Fig. 1.**
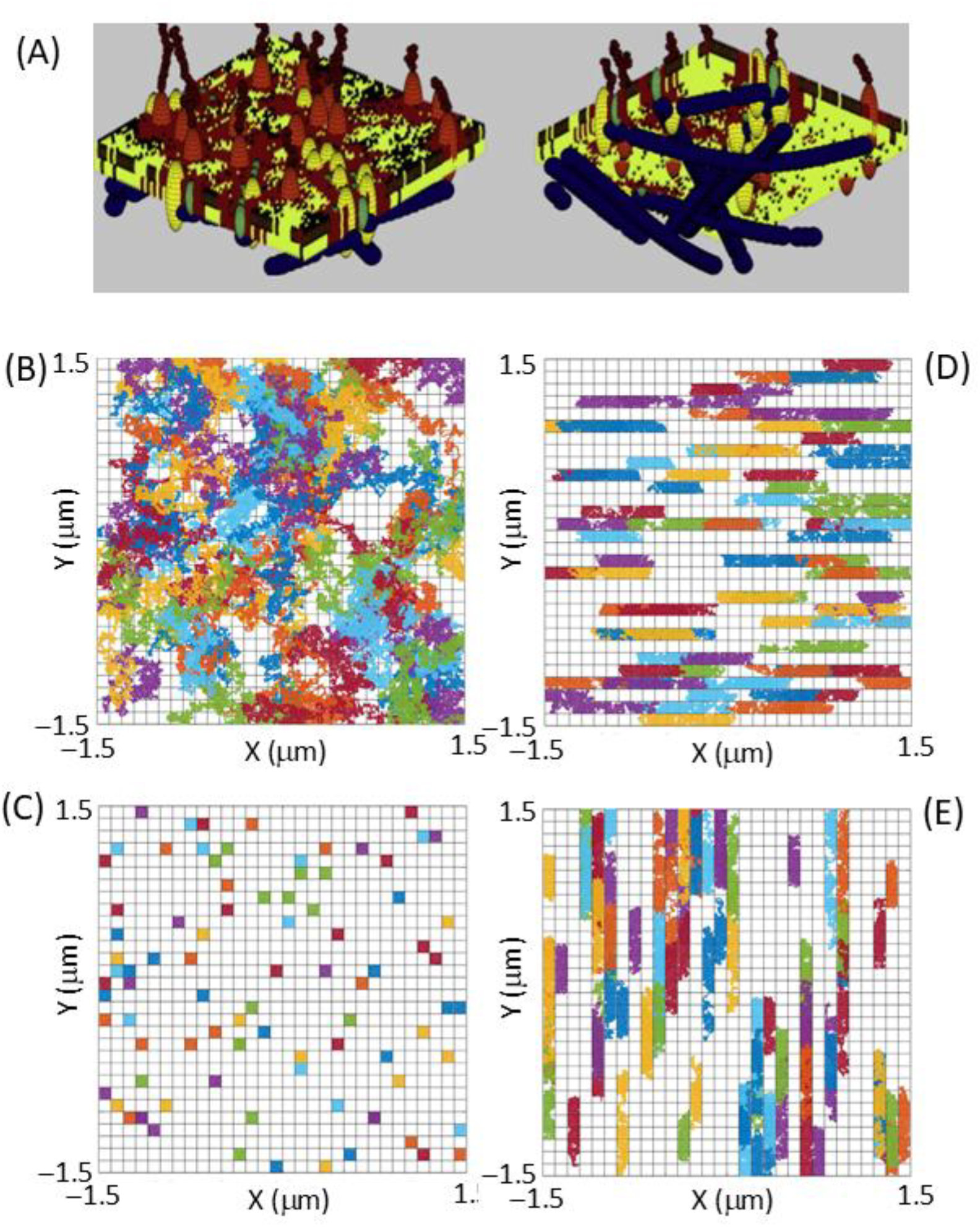
(A) Computer simulation of the interaction between the cortical cytoskeleton and the cell membrane both directly and through integral membrane proteins (Adapted with permission from ‘Hall D. (2010) Effect of heterogeneity on the characterization of cell membrane compartments: I. Uniform size and permeability. Analytical biochemistry 398, 230-244’). (B–D) Simulated diffusion over a 0.1s time period of 90 particles having a D = 1μms^−1^ in a membrane lattice network having an inter-lattice spacing of 100 nm for **(B) Free diffusion** G_X_ = G_Y_ = 1, **(C) Constrained diffusion** G_X_ = G_Y_ = 0, **(D) Vertically constrained diffusion** G_X_ = 1; G_Y_ = 0, and **(E) Horizontally constrained diffusion** G_X_ = 0; G_Y_ = 1.

The vast majority of quantitative experimental and theoretical studies of compartmentalized hop diffusion have considered the compartment properties to be effectively symmetric with regard to their barrier properties **[Kusumi et al. 1993; Sako and Kusumi, 1994; (review Saxton and Jacobson, 1997); Fujiwara et al. 2002; Ritchie et al. 2003; Murase et al. 2004; Ritchie et al. 2005; Wieser et al. 2007; Di Rienzo et al. 2013; Andrade et al. 2015; Goiko et al 2016; Sadegh et al. 2017; de Wit et al, 2018; Mashanov et al. 2021; van der Linden, 2022]**. However, there are some exceptions, notably in a 2011 study of the diffusional characteristics of the CD36 protein, Jaqaman et al. reported the existence of extended linear compartments within the cell membrane of macrophages, with this compartment asymmetry suggested to be caused by the disruption of actin cortical cytoskeleton by microtubule fibers growing close to the cell membrane **[Jaqaman et al. 2011]**. In later work, Di Renzo et al. utilized a two-dimensional pair correlation function analysis of the fluctuations in fluorescence spectroscopy measurements of GFP labelled H-Ras protein to identify diffusive channels in the cell membrane of CHO-K1 cells **[Di Rienzo et al. 2016]**. As discussed by the authors of those works, the existence of such asymmetric channel structures holds potentially important consequences for regulating the interaction of membrane components **[Jaqaman et al. 2011; Di Rienzo et al. 2016]**. In this short letter we develop approximate numerical and analytical methods for simulating and analyzing the Brownian motion of particles within an asymmetrically compartmentalized cell membrane. We then use these quantitative tools to provide comment on the biological advantages offered by such asymmetric arrangements of compartments that relate to modulation of intra-membrane reaction rates and reaction specificity.

## Theory development and results

### Numerical and analytical descriptions of hop diffusion

Thermally driven random movement of the molecular components within a two-dimensional fluid was modelled using an overdamped Langevin formulation of single particle Brownian motion utilizing the particle’s short time diffusion constant, D_0_ **[Hall and Hoshino, 2010; Erban 2014; Zembrzycki et al 2023]** (**Eqn. 1a)**. Using this approach the updated particle position, **r**_p_, at a particular time, t, can be evaluated over an incremental time step, Δt, by solution of Eqn.1a through selection of a random direction α(t) (**Eqn. 1b**) **[Lemons and Gythiel, 1997; Hall and Hoshino, 2010]**.

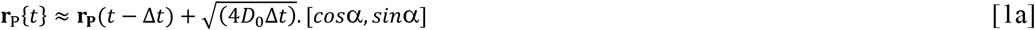

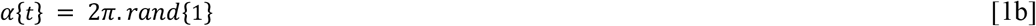

Barriers to diffusive motion were modelled by defining both the linear densities (units of m^−1^) of obstacles in the horizontal, ρ_x_, and vertical, ρ_y_, directions and the obstacle crossing transition probabilities (numbers between 0 and 1) for the horizontal, G_x_, and vertical, G_y_, directions. Assessment of barrier crossing success was based on the trial of a randomly generated number against the set transition probabilities **[Press et al. 2002]**. If the random number challenge criteria was not met the particle was considered to remain at its original position **(Fig. 1) [Saxton, 1990, 1995; Saxton and Jacobson, 1997; Ritchie et al. 2003, 2005; Hall, 2010; Hall and Hoshino, 2010; Rajani et al. 2011]**.

Existing analytical theory has been shown to be capable of describing the effect on diffusive motion by an infinite array of partially permeable barriers **[Kolinski et al. 1986; Powles et al. 1992; cf. Rajani et al. 2011; Weigel et al. 2012; Slezak and Burov, 2021]**. With regard to such hindered random motion occurring in a plane, the governing equations for a particle’s mean squared displacement as a function of sampling interval are described by Eqn. 2a. These equations are defined in terms of the inter-obstacle distance spacing, L_i_, (where i defines either the x or y dimensions and L_i_ = 1/ρ_i_) and a partition constant for barrier crossing in the x and y dimensions, Q_i_, having units of m^−1^ and defined within the range bounded by [0,∞]. The relationship between the direction specific value of the partition constant, Q_i_, and the obstacle transition probability, G_i_, has been previously given and is shown as Eqn. 2b **[Kolinski et al. 1986; Powles et al. 1992; Hall, 2010]**.

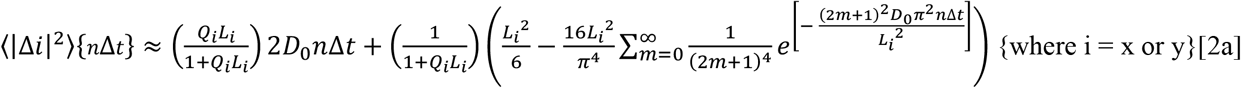

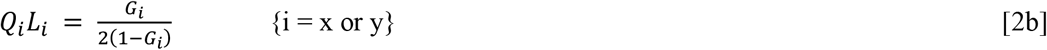

Values of the mean squared displacement in each dimension produced by Eqn. 2a were used to calculate estimates of the time and directionally dependent diffusion constants D_x_{Δt} and D_y_{Δt} **(Eqn. 3a)**. As discussed previously, at a sufficiently high density of barriers these time dependent diffusion constants can be used to evaluate a two-dimensional diffusional probability density function **(Eqn. 3b) [Powles et al. 1992]**.

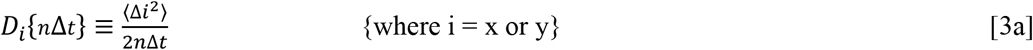

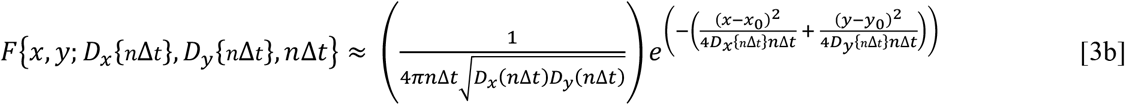

The general soundness of the approximation developed in Eqn. 3is established by comparison of the analytical (Eqn. 3b) and numerical (Eqn. 1) descriptions of both isotropic and anisotropic diffusion caused by non-equal barrier transfer probabilities in the X and Y direction **(Fig. 2)**.

**Fig. 2.**
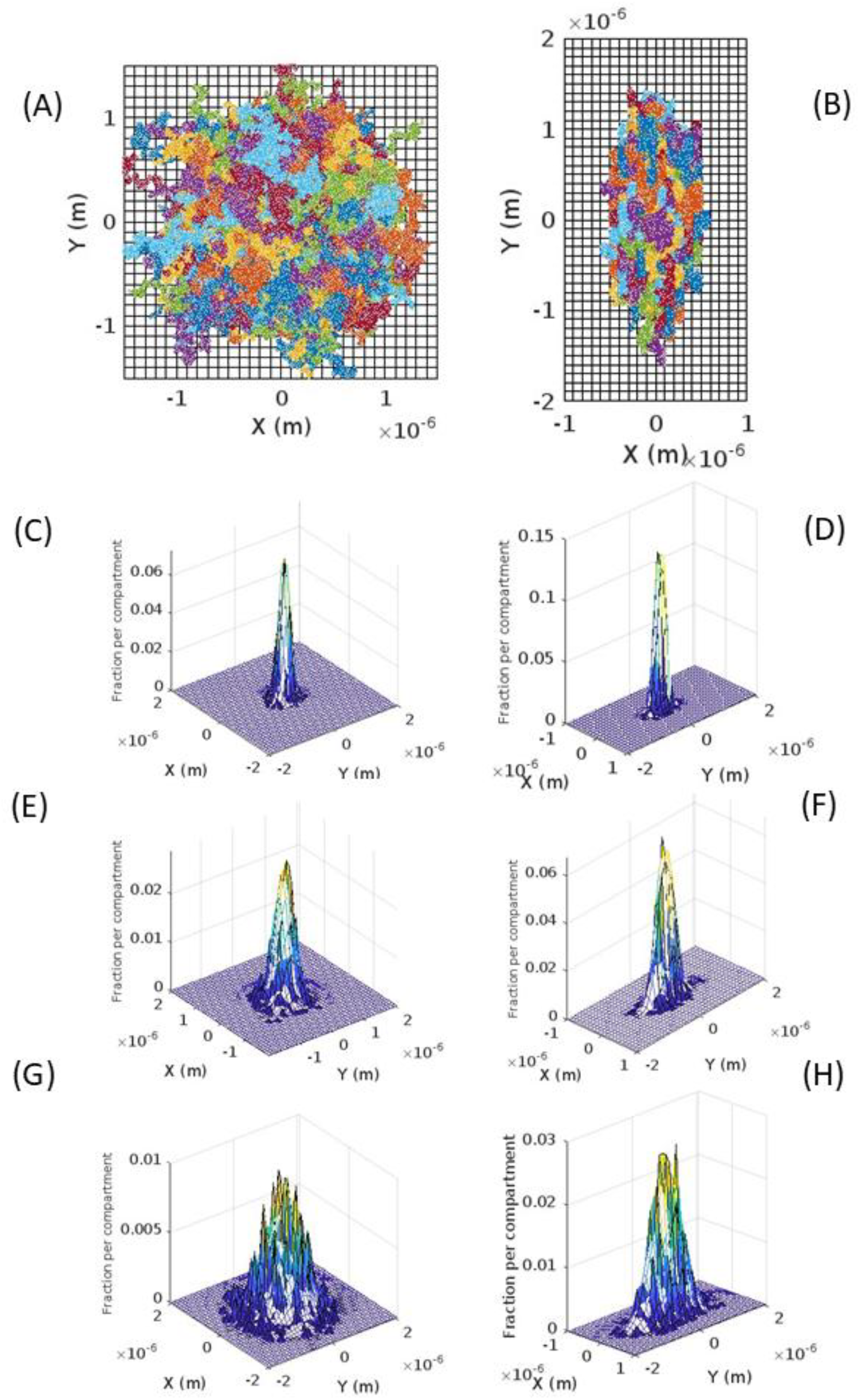
Compartmentalized membrane diffusion as generated by simulation of single molecule trajectories (Eqn. 1) or via integration of the two-dimensional diffusion probability density function (Eqn. 3) (for a particle with D = 1×10^−12^m^2^s^−1^. **(A) Isotropic diffusion:** Overlay of 1000 simulated single molecule trajectories corresponding to free diffusion (i.e. G_X_ = G_Y_ = 1) over a total time of 0.1s. **(B) Anisotropic diffusion:** Trajectory overlay of 1000 simulations of Brownian motion within an asymmetric distribution of barrier probabilities (G_X_ = 0.1, G_Y_ = 1) for 0.1 s. **(C-E)** Two-dimensional histogram of trajectories for isotropic case vs. surface manifold of integrated probability density function calculated using D_X_ = D_Y_ = 1×10^−12^m^2^s^−1^. **(F-H)** Two-dimensional histogram of trajectories for anisotropic case vs. probability manifold calculated using estimates for D_X_ and D_Y_ derived from Eqn. 2 for each time. For (C, D) t= 10ms; for (E, F) t = 30ms and for (G, H) t = 100ms.

### Asymmetric hop diffusion as a means for enhancing reaction rate

One potential advantage afforded to a living cell by a cell membrane featuring an asymmetric distribution of barrier partition values (or an asymmetry in barrier densities) relates to the potential for enhancement of reaction rate between components co-located within a preferred diffusion channel. To provide quantitative insight into the likely degree of rate enhancement, single particle simulations, of the type prescribed by Eqn. 1, were performed to estimate the time required, Δt_RL_, for formation of an encounter complex between a membrane diffusible ligand, L, initially located at a source point, (x_o_, y_o_), and a stationary protein membrane receptor, R, located at point (x_R_,y_R_) at a distance d_12_ from the origin of the diffusing ligand **(Fig. 3a)**. This single particle-based viewpoint of the rate enhancement as a function of initial separation distance is shown by the distribution of receptor ligand encounter times from the simulation of one thousand total particles (i.e. N_TOT_ = 1000) **(Fig. 3b)**. From the subset of diffusible ligand trajectories successfully arriving at the receptor location (blue square) we calculate the average time taken, <Δt_RL_>, and this average is presented in histogram form as a function of barrier asymmetry and initial separation distance **(Fig. 3c)**. We note that, within the limited observation time of 0.1s, there is little (if any) difference in the average encounter time as a function of barrier asymmetry (we will address this, perhaps initially surprising observation, further within the discussion section). The fraction, f_RL_, of individual first encounters between ligand and receptor (occurring within the observation time of 0.1 s, also denoted as f_RL_{0→0.1s}) is presented as a function of the separation distance d_12_ and the barrier asymmetry (described by G_x_ with G_y_ being set equal to 1) **(Fig. 3e)**. We note that the general effect of increased asymmetry in barrier permeability along the orthogonal direction to the target point is to increase the number of successful ligand receptor encounters occurring within the diffusion channel over the observation time of 0.1s. To simplify the process of simulation and analysis presented in Figs. 3a, b, c and e the developed probability density function (Eqn. 3b) was used to determine the probability of occupation of at the grid position x_R_, y_R_ at each timepoint, *B*{*x*_*R*_, *y*_*R*_, *j*Δ*t*} (Eqn. 4a). From this general occupation probability the particular probability of a particle’s first arrival at a time, nΔt, was evaluated as per Eqn. 4b and c with the basic ansatz as follows. Particles occupying the grid position x_R_, y_R_ at each timepoint, *B*{*x*_*R*_, *y*_*R*_, *j*Δ*t*}, are classified into one of the following three types, (i) particles that have newly (first) arrived, (ii) particles that have previously arrived (and subsequently remained at that position), and (iii) particles that have returned to the position (after having made departure from a previous arrival). An approximate, yet surprisingly accurate (as shown by comparison with numerical simulations cf. Figs. 3c vs. 3d and 3e vs. 3f) means for determining the probability of first arrival, *P*_1_*st*{*x*_*R*_, *y*_*R*_, *n*Δ*t*} was developed by correcting the total occupation probability *B*{*x*_*R*_, *y*_*R*_, *j*Δ*t*} for these two latter types of contribution (i.e. those particles having made a previous arrival (with their subsequent remaining at that position), and those particles returning to that position (after having departed from a previous arrival)) using a recursive process involving parameter, M{x_R_, y_R_, nΔt} (Eqn. 4b and c – derivation provided in Appendix 2). Knowledge of the probability function describing a particle’s first arrival time (Eqn. 4a-c) allowed for direct analytical evaluation of the analogues of <Δt_RL_> (**Eqn. 4d**) and *f*_*RL*_{0 → *n*Δ*t*}(**Eqn. 4e**).

**Fig. 3.**
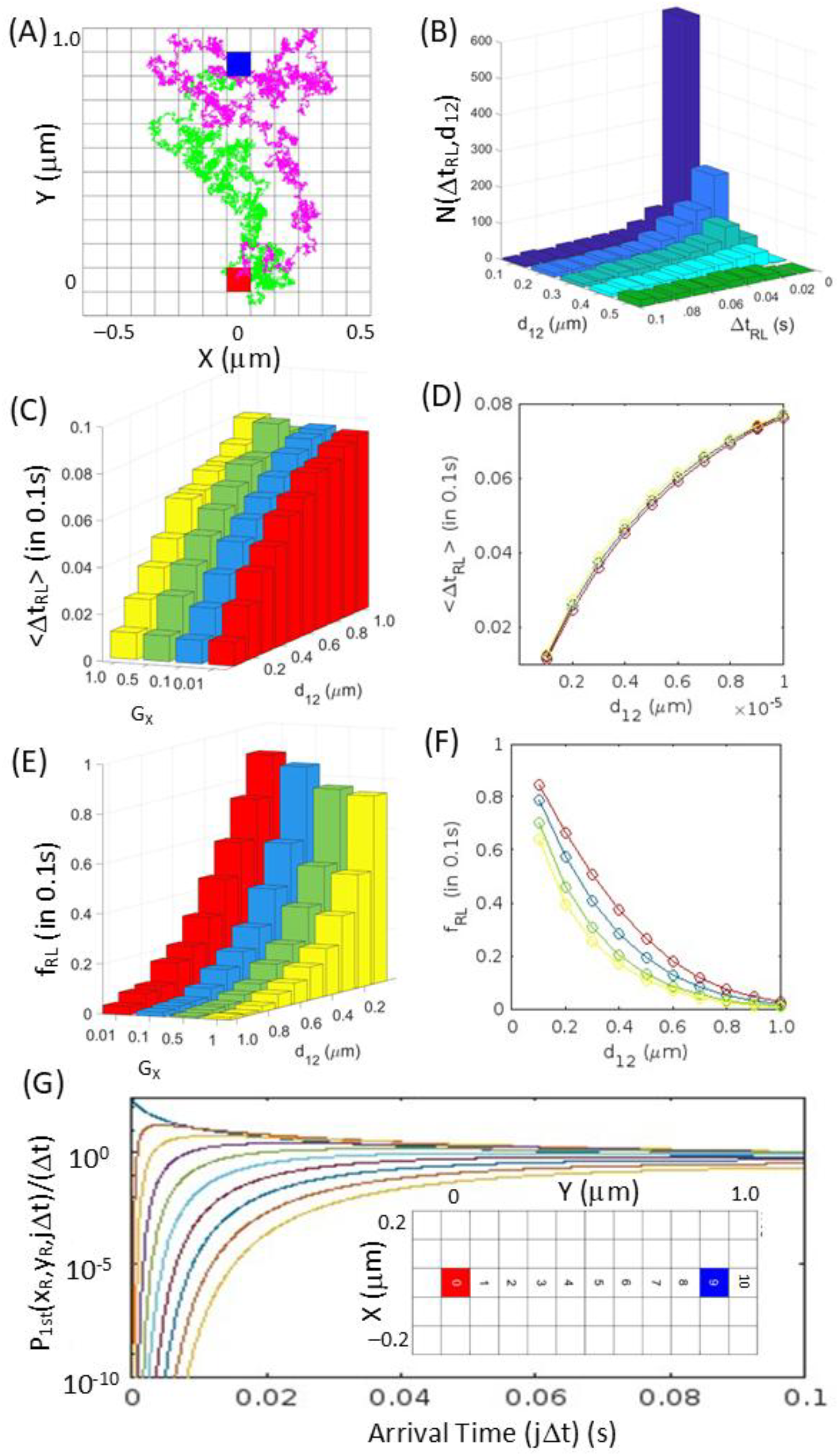
Receptor ligand (RL) encounter rate properties determined by single molecule simulations and application of developed analytical theory. **(A) Encounter trajectory:** Two examples of single molecule simulations of a successful ligand receptor encounter trajectory describing motion of a diffusible ligand (L) released from a source point (x_o_,y_o_ - red square) that is capable of random migration to a stationary membrane receptor (R) located at a more distant point (x_R_,y_R_ - blue square) with these two points separated by a distance d_12_. The time required for each individual random encounter is defined as Δt_RL_ (simulation generated for the free diffusion case: compartment barriers G_X_ = G_Y_ = 1); D_0_ =1×10^−12^ m^2^s^−1^. **(B) Histogram of recorded ligand receptor encounter times made from single particle simulations:** Distribution of Δt_RL_, for ligand/receptor interactions occurring within 0.1s from 1000 trial single particle simulations each with initial separation distances, d_12_ (example demonstrated for free diffusion case: compartment barriers G_X_ = G_Y_ = 1). **(C) Bar chart showing the average time of successful ligand receptor encounters <Δt**_**RL**_**> gained from analysis of single particle simulations:** No significant difference or trend in the average ligand receptor encounter time is observed as a function of barrier permeability asymmetry (G_X_ varied, G_Y_ = 1) at any of the initial separation distances. Analysis of successful encounters occuring within 0.1s evaluated from 1000 trial simulations. **(D) Calculation of average time of successful ligand receptor encounters <Δt**_**RL**_**> using the developed analytical theory (Eqn. 4d):** Average ligand receptor encounter time as a function of barrier permeability asymmetry (G_X_ varied, G_Y_ = 1) for each of the possible initial separation distances (colours define barrier permeabilities as per Fig. 4C). **(E) Bar chart showing the fraction of successful ligand receptor encounters, f**_**RL**_, **evaluated from single particle simulations:** f_RL_ presented as a function of the initial separation distance, d_12_, and the asymmetry of the barrier porosity as shown by G_X_ (with G_Y_ = 1) (note the greater fractional completion extent at lower values of G_X_). Analysis of total trajectories in 1000 trial simulations. **(F) Calculation of the fraction of successful ligand receptor encounters, f**_**RL**_, **using the developed analytical theory (Eqn. 4e):** Average ligand receptor encounter time as a function of barrier permeability asymmetry (G_X_ varied, G_Y_ = 1) for each of the possible initial separation distances (colors define barrier permeabilities as per Fig. 4E). **(G) Probability density function (transformation of Eqn. 4b) reflecting first arrival time, P**_**1**_ ^**st**^**(x**,**y**,**t)/Δt, for diffusion of a particle from ligand source (red square) to the receptor position (blue square), plotted against time:** Each of the ten colored distributions correspond to a different distance of separation, d_12_, from 0.1 to 1μm numbered square in the channel (example given for isotropic case i.e. G_X_ = 1.0; G_Y_ = 1.0). Evaluation of the probability of achieving a first arrival time, P ^st^(x,y,t), can be gotten from the graph by integration of the probability density function,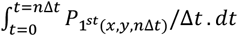.

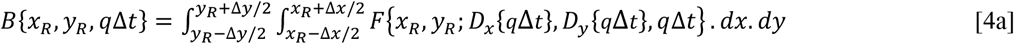

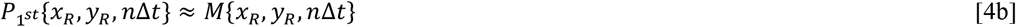

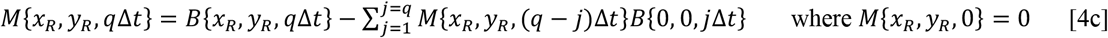

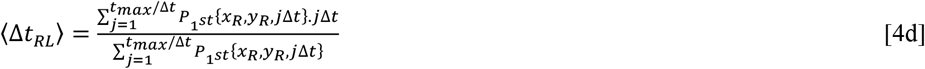

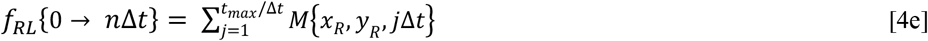

Although it is based on the assumption of diffusion within an unbounded surface, the calculation of the probability of a particle’s first arrival time using Eqn. 4a-c and subsequent quantities derived therefrom, such as <Δt_RL_> using Eqn. 4d **(Fig. 3d)** and *f*_*RL*_{0 → *n*Δ*t*} using Eqn. 4e (**Fig. 3f**), hold the advantage that they are analytical and hence not requiring the application of histogram analysis of the results of numerous trajectory simulations (cf. respectively Fig. 3c and 3e). Examples of the related first arrival probability density functions, *P*_1_*st*{*x*_*R*_, *y*_*R*_, *t*}/(Δ*t*), for ten different separation distances between ligand source and receptor sink are presented for the isotropic case (**Fig. 3g**).

### Asymmetric hop diffusion as a means for enhancing specificity

A second potential advantage afforded to a cell by an asymmetric arrangement of porous membrane compartments is the ability to generate positional specificity in the activation of receptors within the membrane (Fig. 3a). To provide a useful quantitative estimator, we define a positional specificity factor, S, that shows relative enhancement/diminishment of diffusible ligand at a specific position relative to the completely isotropic case (Eqn. 5).

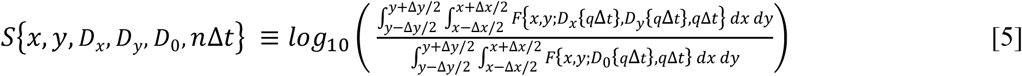

From Fig. 4a-c we see that although the enhancement is modest (up to a factor of 10 for the simulation values adopted here) the extent of negative specificity i.e. exclusion, can differ by many orders of magnitude, making interaction of the diffusible ligand with receptors located in areas outside of the preferred diffusional channel, highly unlikely.

**Fig. 4.**
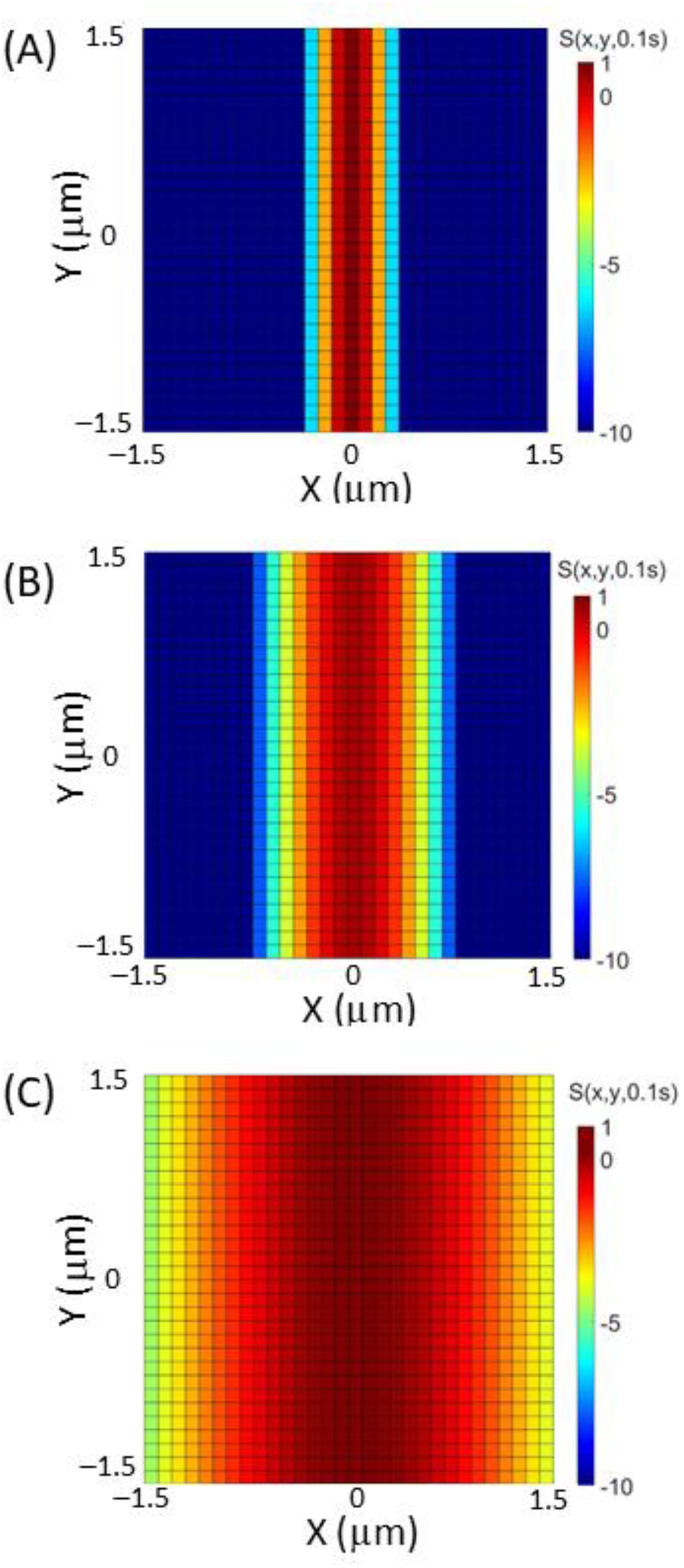
Specific localization of the diffusible ligand as a function of position and asymmetry of barrier permeability. Specificity, S(x,y,t=0.1s) defined by Eqn. 4 for **(A) G**_**x**_ **= 0.01;** (B) G_**x**_ **= 0.1, and (C) G**_**x**_ **= 0.5**. For all cases G_y_ = 1. At zero time the diffusible ligand was located within the central pixel. D_0_ = 1μm^2^s^−1^.

## Discussion

In this paper, methods were developed to simulate and characterize the effects of an asymmetric distribution of barriers, to free movement within the cell membrane. To better relate the theory to experiment all derived theoretical parameters had a directly measurable experimental analogue. The analytical theory was checked against numerical solutions of two-dimensional random motion occurring amongst an array of partially permeable obstacles – with an excellent level of agreement being achieved between numerical results and analytical theory. Outside of the calculation of basic parameters, such as spatially resolved mean squared displacement and diffusion coefficients, analytical expressions were derived for more complex quantities which included, <Δt_RL_>, the average time of a released ligand’s first arrival at its receptor site^2^ (Eqn. 4d), f_RL_{0→nΔt}, the fraction of released ligands achieving a first arrival at their spatially distant receptor site within a finite time interval nΔt (Eqn. 4e), and *S*{*x, y, D*_*x*_, *D*_*y*_, *D*_0_, *n*Δ*t*}, the directional specificity factor describing the ratio of ligand density at a point (x, y), and a specific time, *n*Δ*t*, for the anisotropic versus the isotropic case (Eqn. 5). To provide some perspective on the importance of the current work, before discussing the strengths, limitations and potential ramifications of the model, we first outlay a more detailed history of the study of diffusion within the cell membrane.

### Diffusion in the cell membrane

It is humbling to appreciate that the first quantitative measurements of protein and lipid diffusion in the membrane of living cells were made only fifty years ago^3^ **[Cone, 1973; Poo and Cone, 1974; Peters et al. 1974; Edidin et al. 1976]**. Predominantly carried out using techniques based on optical signal recovery after photobleaching, a number of these early studies revealed very wide distributions of lateral diffusion constants, that ranged from basically 0 μm^2^s^−1^ (immobile fraction) up to ∼10 μm^2^s^−1^ **[Axelrod et al. 1976; Schlessinger et al**., **1976; 1977; Cherry, 1979]**. This diversity in measured values posed a problem for the fluid mosaic model of the cell membrane, which predicted a relatively weak dependence of the lateral diffusion constant on particle size when migrating through continuum liquids having the bulk viscosity of phospholipid bilayers **[Singer and Nicolson, 1972; Saffman and Delbruck, 1975]**. Three general explanations were proffered to account for the large measured range of cell membrane diffusion constants, (i) phase separation of the cell membrane components **[Shimshick and McConnell, 1973; Ohnishi and Ito, 1974; Jacobson and Papahadjopolous, 1975; Helenius and Simons, 1974 (see last section of this review)]**, (ii) frustrated interaction of free membrane components with those components made immobile through their interaction with the relatively stationary fibers of the cytoskeleton and extracellular matrix **[Edelman, 1976; Sheetz et al. 1980; Koppel et al. 1981]**, and (iii) size-dependent density effects induced by two-dimensional crowding within the membrane [**Jacobson et al. 1987; Minton, 1989**]. When single particle observation of membrane located lipids and proteins became technically possible [**Webb, 1977; Geerts et al. 1987; de Brabander et al. 1991; Kusumi et al. 1993; Schmidt et al. 1995; Wawrezinieck et al. 2005**] a second peculiar feature of diffusion within the biological cell membrane became apparent – namely that individual diffusion coefficients calculated from a single particle’s trajectory were often found to be non-Fickian or ‘anomalous’ in nature [**Qian et al. 1991; Saxton and Jacobson, 1997**]. As described by Qian et al. [**Qian et al. 1991; Owen et al. 2009**], anomalous diffusion could be usefully empirically characterized by parameterizing the proportionality between the particle’s recorded mean squared displacement (defined from the positional vector **r**_**p**_ = [x, y]) and the change in observation time interval, nΔt, in terms of an exponent β (**Eqn. 6**).

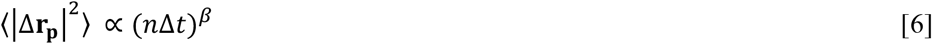

Based on the value of the exponent three general behavioral regimes were defined, (i) Super-diffusive regime β > 1, (ii) Diffusive regime, β = 1, and (iii) Sub-diffusive regime, β < 1, with the final sub-diffusive regime being the case most commonly encountered in experimental investigations^4^ [**Qian et al. 1991; Saxton and Jacobson, 1997**]. Several explanations for sub-diffusion have been advanced with those relating to either phase separation of the cell membrane or frustrated interaction of the lipid/protein membrane components with the cytoskeleton and extracellular matrix both being congruent with experiments of changed diffusion behavior resulting from membrane cholesterol depletion and cytoskeleton/ ECM detachment **[Murase et al. 2004; Vrljic et al. 2005; Sadegh et al. 2017]**. Adopting a phenomenological viewpoint allows for a generalized discussion of the theoretical and biological basis of the effects of barriers to two-dimensional diffusion within the membrane without necessarily assigning an exact biochemical cause^5^ (**see App. Fig. 1**).

### Evidence for anisotropic diffusion in the cell membrane

Whilst numerous investigations have strongly indicated the compartmentalized nature of diffusion of lipids and proteins within the cell membrane **[Kusumi et al. 1993; Sako and Kusumi, 1994; (review Saxton and Jacobson, 1997); Fujiwara et al. 2002; Ritchie et al. 2003; Murase et al. 2004; Ritchie et al. 2005; Wieser et al. 2007; Di Rienzo et al. 2013; Andrade et al. 2015; Goiko et al 2016; Sadegh et al. 2017; de Wit et al, 2018; Mashanov et al. 2021; van der Linden, 2022]** relatively few have explored the potential for these compartments to act as directors of anisotropic motion. Of particular relevance to the current work are three papers (i) the original observations of anisotropic motion of membrane adsorbed concanavalin A in mouse fibroblasts **[Smith et al. 1979]**, (ii) the phenomenological discovery of the asymmetric diffusion of the CD36 membrane protein receptors on the surface of primary human macrophages [**Jaqaman et al. 2011**], and (iii) the technical developments, involving the formulation of a diffusion tensor for describing the diffusional anisotropy of H-Ras-GFP fusion protein within the basal cell membrane of a CHO-K1 cell **[Di Rienzo et al. 2016]**. With respect to this subset of three papers, the initial work by Smith et al. used the fluorescence recovery after photobleaching (FRAP) technique in conjunction with a spatial diffusion tensor formulation to observe the anisotropic movement of fluorescently labelled concanavalin A in a direction parallel to membrane located actin microfilament stress fibers, with an approximately 3-to-4 fold increase in the measured diffusion coefficient parallel vs orthogonal to the stress fiber axis **[Smith et al. 1979]**. Although it was not based on single molecule observation, this work advanced the concept of anisotropic movement and suggested that it was directed by oriented obstacles located within the membrane **[Smith et al. 1979]**. In the work by Jaqaman et al. (**2011**) it was found that microtubules existing proximally to the inner face of the cell membrane created large linear regions of the membrane having no direct/intimate interaction with the cortical actin cytoskeleton. CD36 membrane receptors existing within these extended linear subregions were shown by single particle tracking to exhibit significant diffusional anisotropy along the region covered by the proximal microtubule **[Jaqaman et al. 2011]**. As noted by the authors, such coupling of diffusional anisotropy with compartmentalization offered significant enhancement of both rapidity and sensitivity of CD36 receptor function which is determined by its membrane-based oligomerization **[Jaqaman et al. 2011]**. The work by Di Renzo et al. (**2016**) employed fluorescence correlation spectroscopy in a paired correlation format, simultaneously interrogating neighboring regions within the plane of the cell membrane to determine any potential directional bias in the emergence and entry of fluorescence fluctuations (indicating a preferred movement of the H-Ras-GFP fusion protein) **[Di Rienzo et al. 2016]**. Using this technique the authors could infer significant anisotropy of H-Ras motion in directions aligning with actin fiber mediated cell membrane focal adhesions, thereby suggesting a direct means for cortical actin fiber creation of asymmetric diffusion channels **[Di Rienzo et al. 2016]**.

### Theory developed within the current work for describing asymmetric hop diffusion

A foundation for checking the theory developed within this paper was first laid down using numerical simulations of ligand diffusion within a partially permeable compartmentalized array (Eqns. 1-3) (Figs. 1 – 3). The construction of a closed theory began by incorporating already developed means for describing the mean squared displacement of particles within an infinite array of partially permeable barriers **[Kolinski et al. 1986; Powles et al. 1992;** for alternative solutions see also **Rajani et al. 2011; Weigel et al. 2012; Slezak and Burov, 2021]**, into an analytical solution of the two-dimensional diffusion equation (Eqns. 2 and 3). At each time point, direct evaluation of this 2D-diffusion equation provides a solution that is essentially exact, with any inaccuracies arising from the numerical technique used for its spatial integration (Eqn. 4a). Once formulated Eqn. 4a was used to derive three descriptive parameters relating to the mean first arrival time of ligand receptor encounter, <Δt_RL_> (Eqn. 4d), the fraction of successful encounters within a set time period, f_RL_{0→nΔt} (Eqn. 4e), and a directional specificity factor describing relative spatiotemporal effects of asymmetric compartmentalization on spatial ligand density, *S*{*x, y, D*_*x*_, *D*_*y*_, *D*_0_, *n*Δ*t*} (Eqn. 5). The correctness of these derived parameters was checked against histogram analysis of the results of numerous single particle simulations [Eqn. 1]. Due to the physical insight provided by each of the derived parameters we now provide a short discussion of each in turn.

The average first arrival time of receptor ligand encounter, <Δt_RL_> (Eqn. 4d), represents the mean of the recorded first encounter times between mobile ligands and their stationary receptors, initially spatially separated by a distance, d_12_, under the caveat that this first collision is required to occur within the set maximum allowed time of the simulation. This latter qualifier (i.e. the encounter occurs within the time [0,nΔt]), means that <Δt_RL_> will generally increase with increasing total simulation time and is therefore not a fixed property of the system (as can also be appreciated from the required renormalization term shown in Fig. 4d). As demonstrated by Fig. 3B, the total number of successful encounters within a set time period, will decrease with increasing separation distance. The average first arrival time <Δt_RL_> is an example of a first passage quantity that are typically evaluated by either analysis of numerous numerical simulations **[Rajani et al. 2011]** or by analytical solution of a time dependent master equation [**Iyer-Biswas and Zilman, 2016; Redner, 2023**]. The approach used in this paper, utilized an approximation that greatly simplified the mathematics and yet produced near identical results to the numerical solutions (for a full derivation of Eqn. Set 4 see **Appendix 2**). In brief, this approximation involved consideration of all particles occupying the grid position x_R_, y_R_ at a particular timepoint, *B*{*x*_*R*_, *y*_*R*_, *j*Δ*t*}, as comporting to one of three classes (i) those newly (first) arrived, (ii) those previously arrived/remaining, and (iii) those returning after a prior departure. This explicit treatment of any newly arrived particles at the receptor location x_R_, y_R_, at each time increment, as a distinct class of ligand, allowed for a separate diffusion equation to be initiated and subsequently solved using a starting time set at jΔt. At any subsequent later time, qΔt, the amount of ligand considered as either remaining or previously arrived was jointly evaluated as the sum of the products of the remaining amounts of each ‘newly arrived’ ligand from each prior timepoint jΔt, M(x_R_, y_R_, jΔt), and a normalized 2D diffusion equation evaluated at a virtual origin and a virtual elapsed time calculated from each of the respective ‘zero’ times (i.e. B(x_R_ − x_R_, y_R_ − y_R_, (q−j)Δt) (see Eqn. 4c). Although Eqn. 4c does not explicitly include separate terms to account for particles that are remaining or returning after a prior departure it well describes the sum of these two terms due to the fact that the solution of the density at the virtual origin includes particles that have both not yet left the origin as well as particles that have left and returned. Aside from the relative simplicity of its formulation, the developed approach also offers significant advantages when calculating first arrival times for irregular geometries e.g. irregular geometries of arbitrary angle of presentation. Indeed, although in the current work, both source and sink were considered as square regions, no significant difficulties would be faced for evaluation of average first arrival times for more exotic shapes on the proviso that they be broken down into a series of connected (or unconnected) squares.

The fraction of successful encounters within a set time period, f_RL_{0→nΔt} was evaluated by straight summation of the probability of a particle’s first arrival at each sub-time interval (Eqn. 4e). On the proviso that the approximate technique developed in this paper for calculation of first arrival probability is correct (note that the agreement between the numerical results and the analytical theory is remarkably good cf. Fig. 3E vs 3F), then unlike for the quantity <Δt_RL_>, evaluation of f_RL_{0→nΔt} needs no additional renormalization (as it represents the fraction of all ligands). However, similar to <Δt_RL_>, f_RL_{0→nΔt} will also be a time dependent quantity, becoming larger (within the asymptotic limit of 1) with increasing simulation time.

Within this paper, a specificity factor, *S*{*x, y, D*_*x*_, *D*_*y*_, *D*_0_, *n*Δ*t*}, was defined as the base 10 logarithm of the ratio of anisotropic and isotropic diffusion density profiles at some arbitrary time (Eqn. 5). The base 10 aspect of the logarithm allows for easy assessment of either the excess or diminished densities between the two cases (Fig. 4). Due to its flexible nature i.e. no absolute requirement for a particular initial condition starting geometry, the specificity factor could be applied to all spatial measurements of diffusion in the membrane in either a forward looking (exploring the potential effect of anisotropy generated by a postulated asymmetric barrier geometry overlaid on otherwise isotropic behavior), or backward looking (inferring the degree of anisotropy by dividing the recorded case by simulated/expected isotropic behavior) manner. Such an analysis could be used to search for separation zones within the membrane with this separation aspect potentially reflecting an equal or greater biological importance than the converse case of inclusion.

### Biological consequences of asymmetric hop diffusion

Although the development of new methods for evaluating pertinent ligand receptor encounter quantities (such as the first arrival time, the total fraction of successful encounters and the likely directional specificity of the interaction) represents a potential quantitative advance, it is of perhaps equal or greater importance to recognize the biological significance of the generated results, in relation to subject of diffusion of ligands within the cell membrane. With regard to ligand receptor encounter within a channel existing along a particular directional axis, both the numerical and analytical results (Fig. 3C and D) suggest that for the situation of changing orthogonal direction barrier permeability, there will be (i) little to no effect on the average first arrival time^6^, (ii) a large effect on the fraction of successful receptor ligand encounters (Fig. 3E and 3F). Such a situation indicates that the major effect of asymmetric barrier permeability is to effectively mimic an increase in the concentration of ligand. However, with regard to this point, it should be appreciated that the apparent fluctuation in arrival times seen between the average of the numerical results (gained from a collection of single particle simulations) and the determined analytical values (gained from the developed theory), will be a function of the number of simulations i.e. the number of diffusing ligand replicates. To generate the numerical results, we calculated averages from 1000 simulations, and note that averages derived from such high replicate numbers, closely approximate the values produced by the analytical theory. From the biological perspective, the relevant number of diffusible ligands might be in the single digits. As the effect of a high degrees of barrier asymmetry is to keep more diffusible ligands within the channel, it is to be expected that the degree of noise in values of <Δt_RL_> and f_RL_{0→nΔt} calculated when using fewer simulation replicates will increase as the degree of barrier asymmetry is decreased. Although the relationship between noise and barrier asymmetry as a function of simulation number was not specifically investigated within the current paper, we might reasonably infer on the basis of the previous arguments, that a direct consequence of the existence of diffusing channels would be to smooth out/make more predictable, the timings between cause (diffusible ligand release) and effect (subsequent cellular outcome triggered by the encounter event). From an engineering perspective the outcome of barrier asymmetry would thus make any biological machine more regular and less subject to stochasticity arising from a wide distribution of timings between input and output events i.e. the biological machine would become more ‘machine-like’.

As previously noted, the existence of the type of diffusive channels shown in Fig. 1 could promote the collocation and encounter of biological components belonging to a common reaction/signaling circuit **[Jaqaman et al. 2011; Di Rienzo et al. 2016]**^**7**^. However, as shown by Fig. 4, the converse aspect of this local concentration effect may be just as important i.e. that the concentration of ligand outside of the channel region can be massively diminished. The existence of biological mechanisms for generating spatial separation of diffusible components would facilitate the cell membrane’s ability to carry out regio-specific functions – thereby further extending our view of biological transduction mechanisms as being cell position directed [**Kalappurakkal et al. 2020; Jacobson et al. 2019**].

## Conclusions

Despite the tremendous complexity of the biological cell membrane, several scientific consensuses have been reached. One such consensus is that that the motion of lipid and protein components within the membrane are frequently restricted to relatively small compartments (from 50 – 500 nm diameter) from which they may intermittently escape, in this way ‘hopping’ from one compartment to the next **[Jacobson et al. 2019; Mashanov et al. 2021; van der Linden, 2022]**. In this work, I have presented methods for simulating and characterizing the case where compartments exhibit asymmetric barrier permeability – a situation which results in diffusional anisotropy over short distance scales. Such anisotropy has been previously observed experimentally **[Smith et al. 1977; Jaqaman et al. 2011; Di Rienzio et al. 2016]** and so the situation is undoubtedly biologically relevant, but exactly how, is still a subject of speculation. Recent developments in ultra-microscopy procedures further open the door for the study of cell membrane structure and dynamics (such as those made in high-speed atomic force microscopy **[Hall and Foster, 2022; Hall, 2023a, b]**, electron microscopy **[Hall, 2012; Kaplan et al. 2021; Zabeo and Davies, 2022]** and super-resolution microscopy **[Andrade et al. 2015; Lagerholm et al. 2017; Sil et al. 2020; Eggeling et al. 2025]**. It is hoped that the numerical and analytical methods developed within this paper may prove useful in helping to further explore the biological and biophysical basis of the cell-membrane hop diffusion model and potentially help integrate such understanding into higher-order models of cell behavior such as those incorporating cell growth and division **[Hall, 2023c, Hall 2024]**.

## Acknowledgements

I am grateful for helpful conversations with Drs. A. Yurtsever and K. Ngo at the outset of this work. I would like to thank Prof. W. K. Olson for helpful scientific discussions and for comments made on an early draft of this manuscript. Additional thanks to Mr. A. Richard and Ms. J. Ann for a careful reading of the manuscript and for providing suggestions for improvements in the overall clarity of expression. I gratefully acknowledge support from Rutgers, State University of New Jersey, provided in the form of a Visiting Scientist appointment held within the Department of Chemistry and Chemical Biology, Center for Quantitative Biology.

## Conflict of Interest Statement

D.H. reports no conflict of interest. No humans or animals were harmed during the writing of this article.

## Appendix 1

The present work takes the viewpoint that Brownian motion within the cell membrane is frequently compartmentalized in nature, without necessarily defining the causative nature of the compartments. Although both cytoskeleton/ECM membrane interaction and membrane phase separation have both been shown to be play significant roles in influencing diffusion within the cell membrane, they are not necessarily separate from each other, and their relative contributions to Brownian motion within the cell membrane will depend on their operational length-scales in relation to each other, and to the recorded area, sampling time increment and total observation time employed by the measurement technique. As membrane phase separation may actually be mediated via differential interaction of the various phase components with the cytoskeleton/extracellular matrix these two effects may also be acting in synergy. (**App. 1 - >Fig. 1**) [**Simons and Ikonen, 1997; Kusumi et al. 2005; Owen et al. 2009; Day and Kenworthy, 2009; Trimble and Grinstein, 2015; Goiko et al. 2016; Krapf, 2018; Jacobson et al. 2019; Gupta et al. 2020**]

**Fig. A1.**
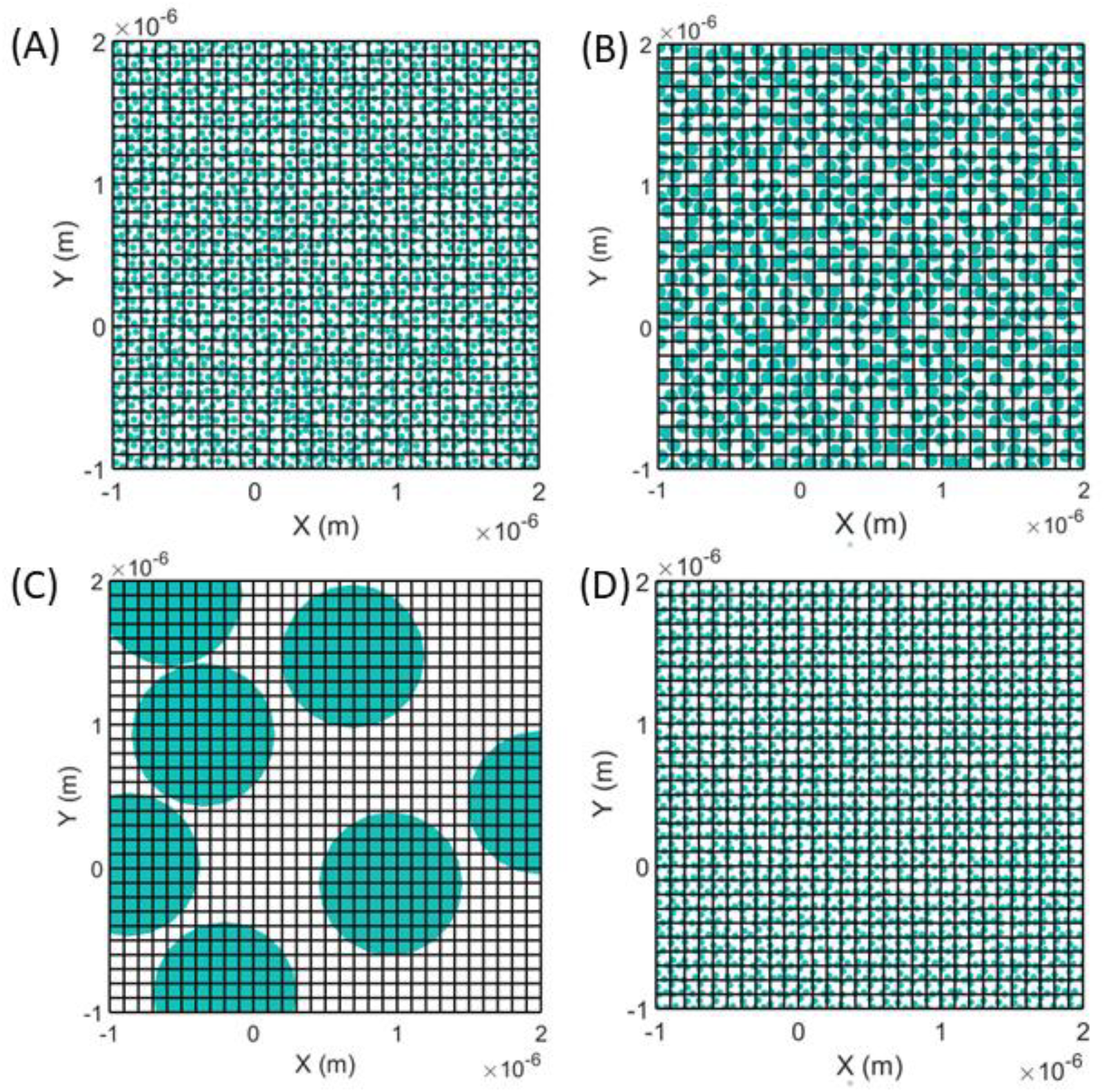
App. 1 Schematic of cell membrane showing barriers caused by either cytoskeleton/extra-cellular matrix (depicted by black lines) or phase separated regions determined by membrane component miscibility (described using cyan circles) Depending on the relative affinity of the diffusing component (whose trajectory is being measured) for either the bulk membrane or the phase separated regions complications can arise in the interpretation of membrane barriers due to the size and association of phase separated regions relative to cytoskeletal/ECM barriers – for example **(A) Phase separated region smaller than fiber induced barriers** (phase regions of 25nm diameter; fiber barriers 100nm spacings)**; (B) Phase separated regions of equivalent size to fiber induced barriers** (phase regions of 50nm diameter; fiber barriers 100nm spacings) **(C) Phase separated regions larger than fiber induced barriers** (phase regions of 500nm diameter; fiber barriers 100nm spacings). (D) **Phase separated region having a strong affinity for fiber induced barrier components** (phase regions of 25nm diameter are preferentially aligned with fiber induced barriers). In all cases the fractional coverage of the surface by the phase separated region is 0.45.

## Appendix 2

To derive the recursive relation shown in Eqn. 4b and c of the main text, one must evaluate the concentration of ligand that is newly arrived to position x_R_, y_R_ within each j^th^ time increment, jΔt. As justification two equation sets are presented as **App. 2 Table 1** and **App. 2 Table 2** that describe the evaluation of this recursion relation for the first three time increments in terms of the recursive index M(x_R_, y_R_, jΔt) (**App. 2 Table 1**) and the fully expanded form (**App. 2 Table 2**). From analysis of the first few terms one can appreciate that the number of terms quickly expands yet the solution can be solved easily via use of a computer program. The key point to grasp is that the amount of ‘newly arrived’ ligand at a certain position and time point, M(x_R_, y_R_, jΔt), is allowed to decay over time by using its initial extent as the starting amplitude to multiply by a decaying normalized diffusion equation centered at the receptor position with a starting reference time equal to jΔt (a decaying point spread function).

**Table A1.** Recursive relation described by Eqn. 4b and c in expanded form.

| Time Increment ($j\Delta t$) | Newly arrived class of ligand within the $j^{\text{th}}$ time increment = $M(x_R, y_R, j\Delta t)$ |
| --- | --- |
| $0\Delta t$ | $M(x_R, y_R, 0\Delta t) = B(x_R, y_R, 0\Delta t)$ |
| $1\Delta t$ | $M(x_R, y_R, 1\Delta t) = B(x_R, y_R, 1\Delta t) - M(x_R, y_R, 0\Delta t) \cdot B(0,0,1\Delta t)$ |
| $2\Delta t$ | $M(x_R, y_R, 2\Delta t) = B(x_R, y_R, 2\Delta t) - M(x_R, y_R, 1\Delta t) \cdot B(0,0,1\Delta t) - M(x_R, y_R, 0\Delta t) \cdot B(0,0,2\Delta t)$ |
| $3\Delta t$ | $M(x_R, y_R, 3\Delta t) = B(x_R, y_R, 3\Delta t) - M(x_R, y_R, 2\Delta t) \cdot B(0,0,1\Delta t) - M(x_R, y_R, 1\Delta t) \cdot B(0,0,2\Delta t) - M(x_R, y_R, 0\Delta t) \cdot B(0,0,3\Delta t)$ |

**Table A2.** Recursive relation described by Eqn. 4b and c in indexed form.

**Appendix 2 Table 2:** Recursive relation described by Eqn. 4b and c in expanded form.
| Time Increment ($j\Delta t$) | Newly arrived class of ligand within the $j^{\text{th}}$ time increment = $M(x_R, y_R, j\Delta t)$ |
| --- | --- |
| $0\Delta t$ | $M(x_R, y_R, 0\Delta t) = B(x_R, y_R, 0\Delta t)$ |
| $1\Delta t$ | $M(x_R, y_R, 1\Delta t) = B(x_R, y_R, 1\Delta t) - B(x_R, y_R, 0\Delta t) \cdot B(0,0,1\Delta t)$ |
| $2\Delta t$ | $M(x_R, y_R, 2\Delta t) = B(x_R, y_R, 2\Delta t) - \{B(x_R, y_R, 1\Delta t) - B(x_R, y_R, 0\Delta t) \cdot B(0,0,1\Delta t)\} \cdot B(0,0,1\Delta t) - \{B(x_R, y_R, 0\Delta t)\} \cdot B(0,0,2\Delta t)$ |
| $3\Delta t$ | $M(x_R, y_R, 3\Delta t) = B(x_R, y_R, 3\Delta t) - \{B(x_R, y_R, 2\Delta t) - \{B(x_R, y_R, 1\Delta t) - B(x_R, y_R, 0\Delta t) \cdot B(0,0,1\Delta t)\} \cdot B(0,0,1\Delta t) - \{B(x_R, y_R, 0\Delta t)\} \cdot B(0,0,2\Delta t)\} \cdot B(0,0,1\Delta t) - \{B(x_R, y_R, 1\Delta t) - B(x_R, y_R, 0\Delta t) \cdot B(0,0,1\Delta t)\} \cdot B(0,0,2\Delta t) - \{B(x_R, y_R, 0\Delta t)\} \cdot B(0,0,3\Delta t)$ |

## Footnotes

1 Depending on the cell type and its position within a tissue the ECM will be variably composed of fibrous proteins (such as collagen and elastin), fibrous proteoglycans (such as heparin sulfate, chondroitan sulfate and keratan sulfate), and the polysaccharide hyaluronic acid. The ECM is connected to the membrane via specialist proteins known as laminin and fibronectin that are either integral membrane proteins (as in the case of laminin) or linkage proteins (as in the case of fibronectin which binds to the transmembrane protein integrin) **[Theocharis et al. 2016]**.

2 Also known as the mean first passage time **[Iyer-Biswas and Ziolman, 2015]**.

3 At the time of writing this piece (January 2025).

4 ^4^ Super-diffusion implies active transport of a membrane component, a process typified by a non-zero mean displacement, such as might be generated by mutual diffusion effects, action of membrane located motor protein or displacement caused by ‘pushing’ or ‘pulling’ resulting from membrane-associated fiber extension or contraction. The three major types of linear fibers making up the cell cytoskeleton, microfilaments (formed from actin), intermediate filaments, and microtubules (formed from tubulin) all form linearly persistent fibers with the polymerization/depolymerization process frequently under the control of a hydrolysable nucleotide triphosphate [e.g see **Korn et al. 1987; Hall, 2003**].

5 Due to its abstract (mathematical) nature the current formalism can accommodate multiple mechanisms for inducing membrane compartmentalization including those not addressed here **[Jacobson et al. 1987; Minton, 1989; Hall, 2008]**. However perhaps the most easily countenanced is the cytoskeleton/ extracellular matrix induced barrier effect **[Sadegh et al. 2017; Krapf, 2018; Gupta et al. 2020; Kalappurakkal et al. 2020]**.

6 At the level of the simulation, the fact that increased barrier asymmetry does not increase the value of <Δt_RL_> is to be expected, due to the fact that the orthogonal components of the tensor for any true diffusion process must be uncorrelated and hence zero.

There is also the possibility that such extended channels could act as alignment/adsorption foci in disease states that target the cell membrane [Sasahara et al. 2010; Hall and Edskes, 2009]

## Notes

### Competing Interest Statement

The authors have declared no competing interest.

### Funding

- Rutgers, The State University of New Jersey: Visiting Scientist Appointment

